# Machine Learning for Population Genetics: A New Paradigm

**DOI:** 10.1101/206482

**Authors:** Daniel R. Schrider, Andrew D. Kern

## Abstract

As population genomic datasets grow in size, researchers are faced with the daunting task of making sense of a flood of information. To keep pace with this explosion of data, computational methodologies for population genetic inference are rapidly being developed to best utilize genomic sequence data. In this review we discuss a new paradigm that has emerged in computational population genomics: that of supervised machine learning. We review the fundamentals of machine learning, discuss recent applications of supervised machine learning to population genetics that outperform competing methods, and describe promising future directions in this area. Ultimately, we argue that supervised machine learning is an important and underutilized tool that has considerable potential for the world of evolutionary genomics.

---

Population genetics over the past 50 years has been squarely focused on reconciling molecular genetic data with theoretical models that describe patterns of variation produced by a combination of evolutionary forces. This interplay between empiricism and theory means that many advances in the field have come from the introduction of new stochastic population genetic models, often of increasing complexity, that describe how population parameters (e.g. recombination or mutation rates) might generate specific features of genetic polymorphism (e.g. the site frequency spectrum). The goal, broadly stated, is to formulate a model that describes how nature will produce patterns of variation that we observe. With such a model in hand, all one would have to do would be to estimate its parameters, and in so doing learn everything about the evolution of a given population.

Thus an overwhelming majority of population genetics research has focused on classical statistical estimation from a convenient probabilistic model (i.e. the Wright-Fisher model), or through an approximation to that model (i.e. the coalescent). The central assertion here is that the model sufficiently describes the data such that insights into nature can be made through parameter estimation. This mode of analysis that pervades population genetics is what Leo Breiman [1] famously referred to as the “data modeling culture,” wherein independent variables (i.e. the evolutionary and genomic parameters) are fed into a model and the response variables (some aspect of genetic variation) come out the other side. Models are validated in this worldview through the use of goodness-of-fit tests or examination of residuals (a recent modern example can be found in [2]).

In this review we argue that population genetics as a field might turn to a different mode of analysis, that of the “algorithmic modeling culture,” or what is now commonly called machine learning (ML). Over the past decade machine learning methods have revolutionized entire fields, including speech recognition [3], natural language processing [4], image classification [5], and bioinformatics [6, 7]. However, the application of machine learning to problems in population and evolutionary genetics is still in its infancy, save for some pioneering examples [8–17]. Machine learning approaches have a number of desirable features, and perhaps foremost among them is their potential to be agnostic about the process that creates a given dataset. Machine learning, as a field, aims to optimize predictive accuracy of an algorithm rather than perform parameter estimation of a probabilistic model. What this means in practice is that ML methods can teach us something about nature, even if our models used to describe nature are imprecise. An equally important advantage of the machine learning paradigm is that it enables the efficient use of high-dimensional inputs which act as dependent variables, without specific knowledge of the joint probability distribution of these variables. Inputs that consist of thousands of variables (a.k.a *features* in the ML world) have been used with great success (e.g. [18, 19]) and increases in the number of features can often yield greater predictive power [1]. Given the ever-increasing dimensionality of modern genomic data, this is a particularly desirable property of machine learning. In this paper we describe several examples where, through a hybrid of the “data modeling” and “algorithmic modeling” paradigms, machine learning methods can leverage highdimensional data to attain far greater predictive power than competing methods. These early successes demonstrate that ML approaches could have the potential to revolutionize the practice of population genetic data analysis.

## An Introduction to Machine Learning

Machine learning is generally divided into two major categories (though hybrid strategies exist): supervised learning [20] and unsupervised learning [21]. Unsupervised learning is concerned with uncovering structure within a dataset without prior knowledge of how the data are organized (e.g. identifying clusters). A familiar example of unsupervised learning is principal component analysis (PCA), which in the context of population genetics is used for discovering unknown relatedness relationships among individuals. PCA takes as input a matrix of genotypes (often of very high dimensionality) and then produces a lower dimensional summary that can reveal how genotypes cluster. An excellent example of the application of PCA to population genetics can be found in Novembre *et al.* [22] where PCA was used to show how relationships among individuals sampled from Europe largely mirrored geography. Supervised learning, on the other hand, relies on prior knowledge about an example dataset in order to make predictions about new data points. Generally, supervised ML is concerned with predicting the value of a response variable, or *label* (either a categorical or continuous value), on the basis of the input variables/features. Supervised learning accomplishes this feat through the use of a *training set* of labeled data examples, whose true response values are known in order to train the predictor (see Box 1 for more detail).

There have been a multitude of important applications of unsupervised machine learning in evolutionary genomics beyond PCA. One popular methodology that has been wildly successful in population and evolutionary genetics is hidden Markov models (HMMs; see [23]). HMMs are a class of probabilistic graphical model that are well suited to segmenting data that appears as a linear sequence, such a chromosomes. For instance with phylogenetic data, HMMs have been used to uncover differences in evolutionary rates along a chromosome [24, 25]. Furthermore, HMMs have been used to infer how the phylogeny itself changes across chromosomes due to recombination [9, 26, 27]. In the context of population genetic data HMMs have been leveraged to detect regions of the genome under positive or negative selection [10] as well as to localize selective sweeps [11, 28].

Although unsupervised ML has been deployed widely and effectively throughout the field, to date there has been less attention paid to supervised learning. Here we give a brief overview of the paradigm of supervised ML and highlight recent population genetic studies leveraging these approaches.

#### Box 1. Supervised Learning in draft form

Supervised ML approaches algorithmically create from a given dataset a function that takes as input a vector and then emits a predicted value for each data point. More formally, these methods learn a function, *f*, that predicts a response variable, *y*, from a *feature vector*, ***x***, containing *M* input variables, such that: *f*(***x***) = *y*. If *y* is a categorical variable, we refer to the task as a *classification* problem, whereas if *y* is a continuous variable we refer to it as *regression*. In supervised learning, the objective is to optimize *f*: ***x*** → *y* using a “training set” of labeled data (i.e. whose response values are known). That is, we assume we have a set of training data of length *n* of the form {(***x***_**1**_,*y*_1_),…, (***x***_***n***_, *y_n_*)}, where ***x*** ∈ ℝ^*M*^. A variety of learning algorithms exist which can create functions that can perform either classification or regression, including *support vector machines* (SVMs [29]), *decision trees* [30] and *random forests* [31], *boosting* [32], and *artificial neural networks* (ANNs [33]) which in modern form are subsumed under the umbrella of *deep learning* [34]. These algorithms differ in how they structure and train *f* (see brief descriptions in the Glossary).

To proceed with building *f* we must define a *loss function*, *L*, that indicates how good or bad a given prediction is. A simple choice for a loss function in the context of classification would be the indicator function such that *L*(*f*(***x***), *y*) = **1** (*f*(***x***) ≠ *y*). For regression one might consider the squared deviation *L*(*f*(***x***), *y*) = (*f*(***x***) − *y*)^2^. Finally, we define the *risk function*, which is typically the average value of *L* across the training set. “Training” is the process of minimizing this risk function.

Once training is complete, we must evaluate our performance on an independent test set. This step allows one to assess whether *f* has become sensitive to the general characteristics of the problem at hand, rather than characteristics particular to data examples in the training set (what is known as *overfitting*). For *binary classification* we might characterize the *false positive* and *false negative* rates or related measures such as *precision* and *recall*. A particularly helpful construct in the case of multiclass classification is the *confusion matrix*, which is simply the contingency table of true vs. predicted class labels for each class. For regression, one could use any tool for evaluating model fit (e.g. *R*^*2*^) or examine the distribution of values of one or more loss functions. Residuals can also be checked for evidence of bias in order to anticipate which types of data are likely to produce erroneous predictions.

**Figure Box 1.**
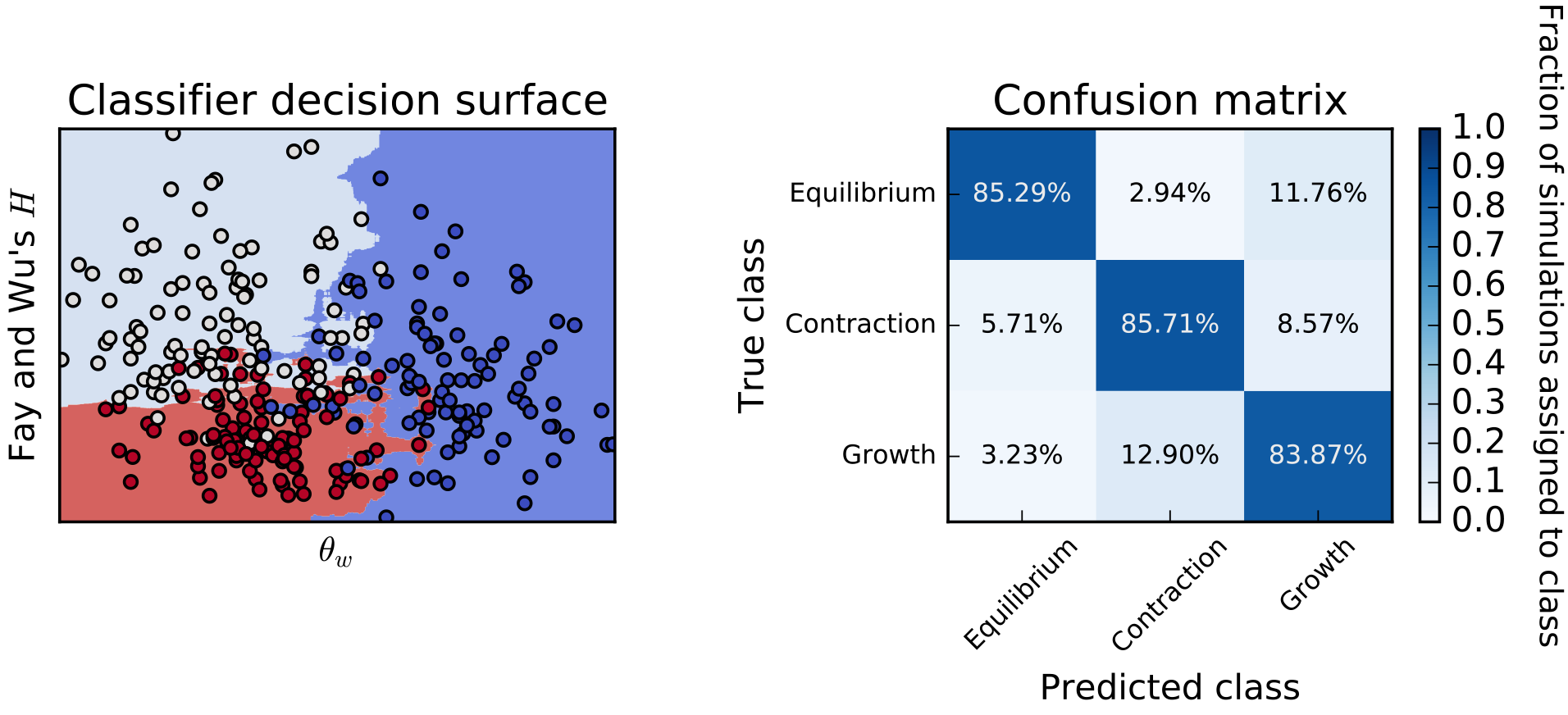
An example application of supervised machine learning to demographic model selection. We simulated population samples experiencing no population size change (Equilibrium), a recent instantaneous population decline (Contraction), or recent instantaneous expansion (Growth). We then trained a variant of a random forest classifier [35], which is an ensemble of semi-randomly generated decision trees, to discriminate between these three models on the basis of a feature vector consisting of two population genetic summary statistics [36, 37]. On the left we show the decision surface: red points represent the Growth scenario, dark blue points represent Equilibrium, and light blue points represent Contraction. The shaded areas in the background show how additional data points would be classified—note the nonlinear decision surface separating these three classes. On the right, we show the confusion matrix obtained from measuring classification accuracy on an independent test set. Data were simulated with Hudson’s ms [38], and classification was performed via scikit-learn [39].

## Why use Machine Learning?

Our basic description of supervised ML approaches in Box 1 demonstrates their central rationale: ML focuses on algorithmically constructed models with optimal prediction as their goal rather than parametric data modeling. Furthermore, ML offers several advantages in addition to accurate prediction. Perhaps most important among them is the ability to circumvent using idealized, parametric models of the data when labeled training data can be obtained from empirical observation (an example of this scenario is given in the following section). Indeed in such cases we can use ML to train algorithms to recognize phenomena as they are in nature, rather than how we choose to represent them in a model. Further, in cases where empirically derived training sets are not available, simulation can be used to generate training sets. This ability to use simulation as a stand-in for observed data is key for population genetics applications, where adequately sized datasets with high-confidence labels are currently hard to obtain. While using simulation for training obviates the model agnosticism that is so attractive about ML, discriminative ML models are more robust to model misspecification than traditional data models [40].

Even when empirical training data cannot feasibly be obtained, there are notable advantages of supervised ML methods. Most importantly, these methods are specifically geared towards using high-dimensional data as input. Typically, classical statistical methods suffer from what has been called the “curse of dimensionality” whereby high-dimensional data become sparse and thus very difficult to fit models to. On the other hand, most supervised ML methods perform better when the input data has a large number of features, in what is commonly called the “blessing of dimensionality” (e.g. [1, 41]). A good example of this comes from the highly cited work of Amit and Geman [18] on using a random forest-like procedure for handwriting recognition: it took as input a feature vector containing thousands of variables, and proved highly accurate. In a more modern setting, *deep learning* methods have been shown both theoretically and in practice to be able to circumvent the curse of dimensionality in many settings [42, 43]. This attribute lends significant strength to population genetics analysis: while inferences are traditionally based on a single summary statistic devised for the given task (e.g. [36, 44–49]), below we describe several recent studies which demonstrate that far greater statistical power can be achieved by simultaneously examining multiple aspects of genetic variation across the genome. Importantly, many ML methods offer direct ways to assess which features of the input are driving inferences, information which can yield insights about the underlying processes [1].

The last benefit we wish to touch upon is computational efficiency. While training of supervised ML algorithms is computationally costly—especially if simulation is used for the training set— once an algorithm is trained, prediction from it is exceedingly fast even in situations where a large number of predictions is required (e.g. genome-wide scans). This means that there will be an upfront cost to training (typically hours or days), but genome-wide inference proceeds rapidly thereafter. Moreover, because many ML approaches (e.g. deep learning) have the ability to generalize beyond their input parameters (e.g. [50]), training sets can be considerably smaller than those used by approaches such as approximate Bayesian computation (ABC [51]; also see Box 3).

## Supervised ML in population genetics by training on real data: finding purifying selection

When empirically derived training data are available, supervised machine learning can be used to make accurate predictions in data sets that cannot be adequately modeled with a reasonable number of parameters. For instance, a current goal in modern genomics is to be able to predict functional regions of the genome using bioinformatics techniques. While there are numerous sources of information to leverage for this problem, including comparative [25] and functional genomics [52], the best manner in which to incorporate population genomic variation to aid in these predictions is a matter of active research. Towards this end we recently used a supervised ML approach to discriminate between genomic regions experiencing purifying selection and those free from selective constraint on the basis of population genomic data alone [15]. In this study we used a support vector machine (SVM) that employed as input the site frequency spectrum (SFS) from all 1,092 individuals from the Phase I release of 1000 Genomes dataset which consisted of 14 population samples from diverse global locations [53]. Had we attempted to use all these data simultaneously in a “classical” population genetics setting we would have been forced to fit a demographic model that described the joint divergence and population size changes of all 14 population samples; a daunting task indeed. While the SFS is well-known to be affected by demography as well as selection [54], by constructing a training set of regions experiencing purifying selection (inferred from a phylogenetic comparison of non-human mammals) we were able to effectively sidestep the intractable problem of modeling the joint demographic history of the dataset. We were then able to both train and test an SVM on empirical data, achieving ~88% accuracy [15].

By comparing the predictions from this classifier, which reveal purifying selection occurring in recent evolutionary history, with phylogenetic signatures of more ancient selection, we were able to identify regions showing evidence of functional turnover in the human genome. We showed that these candidate regions were highly enriched in the regulatory domains of genes important for proper central nervous system development. Moreover, another group [55] recently found that the presence of these candidate regions near a gene was more predictive of human-specific changes of expression in the brain than was the presence of well-known human-accelerated regions (HARs) identified from inter-specific comparisons [56]. This result lends credence both to our own predictions and more generally to the utility of supervised ML approaches in evolutionary genetics.

## Finding selective sweeps in the genome

The population genetic question that has received the most attention from ML approaches is that of detecting selective sweeps: the signature left by an adaptive mutation that rapidly increases in allele frequency until reaching fixation [57]. While the classical population genetic strategy for finding sweeps has been to carefully devise test statistics sensitive to selective perturbations [36, 44–49], in recent years several groups have begun leveraging combinations of statistics through supervised ML to improve inferential power. While each of these methods differ in the exact combination of summary statistics used, their unifying feature is that training sets are generated using coalescent simulations with and without selective sweeps First among these was Pavlidis et al. [12], who used a SVM to combine Kim and Nielsen’s *ω* statistic (which measures the spatial pattern of LD expected around a sweep [47]) with Nielsen et al.’s composite-likelihood ratio (a.k.a. CLR, which highlights the spatial skew in the SFS expected around a sweep [58]). They found that these two statistics in concert had greater power to detect sweeps. Ronen et al. [14] took the approach of encoding the SFS as the feature vector (i.e. each bin in the SFS is one feature), and then used an SVM to discriminate between selective sweeps and neutrality. Lin et al. [8] used *boosting* to identify sweeps on the basis of a feature vector containing six different summary statistics each measured across a number of genomic subwindows surrounding the focal window. In a related effort, Pybus et al. [13] recently used a series of boosting classifiers to detect selective sweeps and classify them according to whether they have reached fixation (complete vs. incomplete) as well as by their timing (recent vs. ancient). Finally, in Schrider and Kern [16], we describe S/HIC, which uses a variant of a *random forest* [31] called an Extra-Trees classifier [35] to detect both classic “hard sweeps” from *de novo* mutations and “soft sweeps” resulting from selection on previously segregating variants [59, 60]. As described in Box 2, S/HIC is able to detect sweeps with high sensitivity and specificity even in the face of nonequilibrium demography, which confounds many other methods. The success of S/HIC and the other efforts listed above demonstrates that an appropriately designed machine learning approach can make rapid advances in performance on difficult problems that have received attention for decades.

#### Box 2 a closer look at S/HIC

We recently introduced a method called S/HIC [16], which uses a feature vector designed to be not only sensitive to hard and soft sweeps, but also robust to the confounding effects of both linked positive selection (i.e. “soft shoulders” [61]) and non-equilibrium demography [54, 62]. This feature vector included values of 9 different statistics that were each measured in a number of adjacent subwindows (Figure Box 2, below), in a similar vein to Lin et al.’s evolBoosting [8]. What set this feature vector apart is that for each statistic, the value in each subwindow was normalized by dividing by the sum across all subwindows. Thus, the true value of a given statistic in a given subwindow is ignored, while the relative values across the larger window are examined. The reasoning behind this choice is that while demographic events may affect values of population genetic summaries genome-wide (which S/HIC ignores), selective sweeps may result in more dramatic localized skews in these statistics (which S/HIC captures). The results of this design are impressive: S/HIC is able to detect sweeps under challenging demographic scenarios, often with no loss in power even when the demographic history is grossly misspecified during training (e.g. if there is an unknown population bottleneck), a scenario which catastrophically compromises many other methods [16, 63]. Thus, ML methods—especially those with appropriately designed feature vectors—can be robust to modeling choices even when training data are simulated.

In Figure Box 2 we illustrate S/HIC’s classification strategy and the values included in its feature vector. This figure demonstrates how much additional information S/HIC utilizes in making its predictions in comparison to more traditional population genetic tests, especially those relying on a single statistic. In particular, the S/HIC feature vector not only includes multiple statistics each of which is designed to capture different aspects of genealogies, but also how these statistics vary along the chromosome. In addition to greater robustness to demography as discussed above, incorporating all of this information yields greater discriminatory power, and for this reason such multidimensional methods will be preferable to univariate approaches. We recently applied S/HIC to six human populations with complex demographic histories, where it revealed that soft sweeps appear to account for the majority of recent adaptive events in humans [64]; the success of this analysis demonstrates the practicality of applying such ML strategies to real data.

**Figure Box 2.**
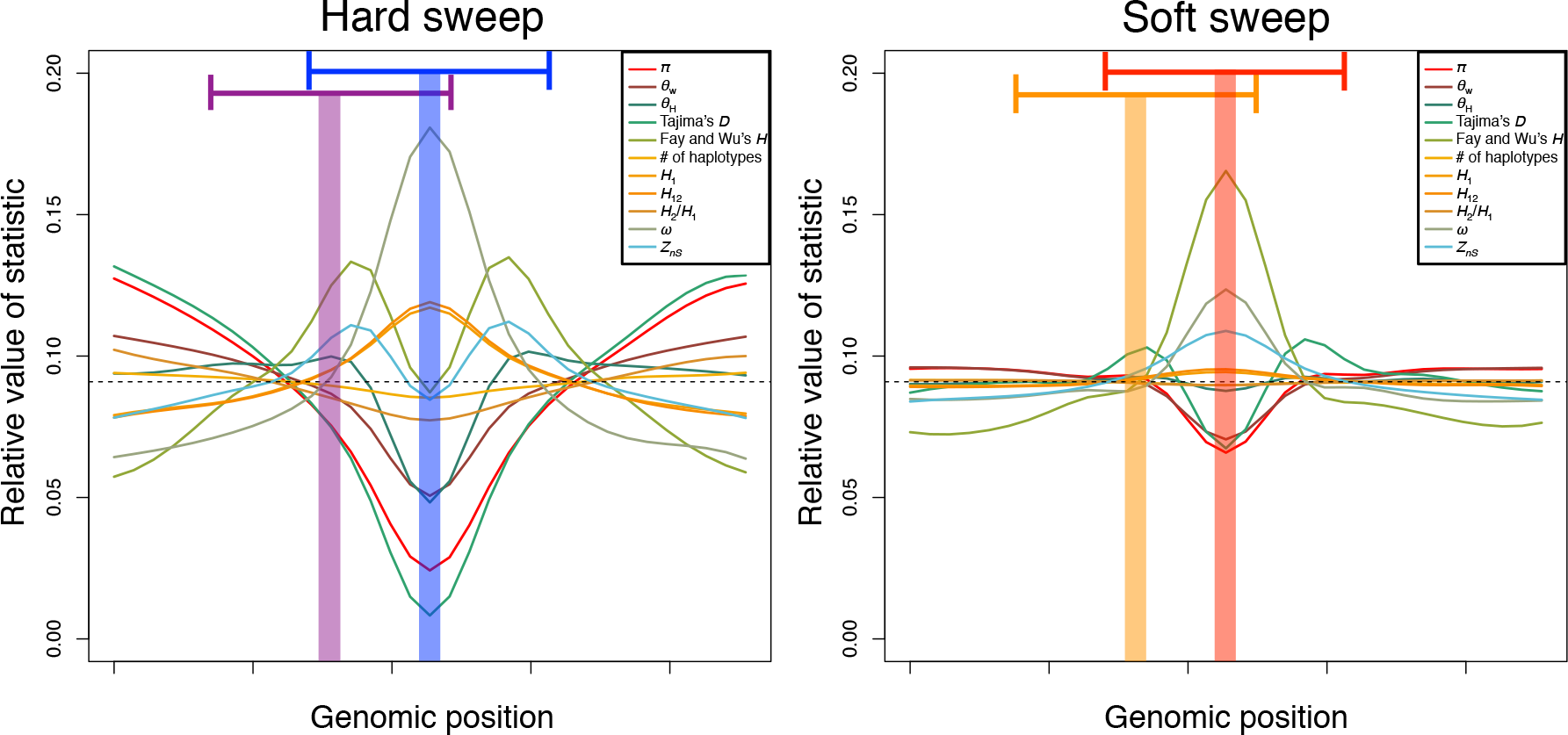
A visualization of S/HIC’s feature vector and classes. The S/HIC feature vector consists of *π* [65], 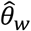 [37], 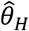 [36], the number of distinct haplotypes, average haplotype homozygosity, *H*_12_ and *H*_2_/*H*_1_ [66, 67], *Z*_*nS*_ [46], and the maximum value of *ω* [47], The expected values of these statistics are shown for genomic regions containing hard and soft sweeps (as estimated from simulated data). Fay and Wu’s *H* [36] and Tajima’s *D* [48] are also shown, though these may be omitted from the vector as they are redundant with *π*, 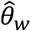 and 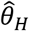. In order to classify a given region, the spatial patterns of these statistics are examined across a genomic window in order to infer whether the center of the window contains a hard selective sweep (blue shaded area on the left, using statistics calculated within the larger blue window), is linked to a hard sweep (purple shaded area and larger window, left), contains a soft sweep (red, on the right), is linked to soft sweep (orange, right), or is evolving neutrally (not shown).

The methods listed above have two commonalities: they use machine learning to perform classification on multidimensional input, and they handily outperform more traditional univariate methods. However, these methods also differ from one another substantially in a number of facets: the particular machine learning framework used, the makeup of the feature vector, and the types of sweeps they seek to detect. Thus, the success of these methods underscores not only the power but also remarkable flexibility of supervised ML. By working within the supervised ML paradigm one can effectively tailor a predictor to whatever task is at hand simply by altering the construction of the feature vector and training dataset, and in so doing, make more detailed predictions than is possible using a single statistic.

Unlike the problem of detecting purifying selection, for which a training set may be constructed, we lack an adequate number of selective sweeps whose parameters are known precisely (e.g. the time of the sweep, strength of selection). Thus, the studies described above used simulation to generate training sets. The general idea is to simulate data from one or a number of population genetic models in which parameters are either specified precisely or defined by prior distributions, use those data to train an ML algorithm, and then perform either classification or regression (i.e. parameter estimation). In this context supervised ML allows for likelihood-free inference of population genetic models similar in spirit to ABC. While, like ABC, this approach requires modeling assumptions, it nonetheless offers numerous advantages as described in Box 3 where we contrast ABC with supervised ML

#### Box 3 Comparing Supervised ML and ABC for population genetic inference

Using supervised ML with training data simulated from a specified set of population genetic models is similar in spirit to approximate Bayesian computation (ABC), save for some notable distinctions. ABC begins by simulating a large number of examples whose model parameters are drawn from prior distributions then summarizes these simulations with vectors of population genetic summary statistics. Next, only those simulations most similar to the observed dataset are retained—a process known as rejection sampling—to approximate the probability distribution for each parameter value given the observed data. ABC is easy to implement, flexible, and has been proven effective in a number of scenarios. However, ABC has some important drawbacks that ML overcomes. Most importantly, when using large feature vectors, ABC is susceptible to the curse of dimensionality [68]—much effort has therefore gone into dimensionality reduction and feature selection for ABC (reviewed in [69]). While this is so, reducing dimensionality might lead to a loss of information if the remaining summaries are not sufficient statistics of the data. This contrasts with modern ML algorithms, which can benefit from high dimensional data, rather that suffer from them.

A second drawback of ABC is its computational burden. While both ML and ABC require a large number of simulations, ABC does not make efficient use of all of this computation because it typically depends on rejection sampling. Work has been done to retain more of the simulations in ABC, for instance by weighing their influence on parameter estimation according to their similarity to the observed data [70]. However ML methods naturally use all of the simulations to learn the mapping of data to parameters. Further, deep learning methods have the potential to generalize non-locally [42], allowing them to make accurate predictions for data quite different from those in the training set. For these reasons, ML may require considerably fewer simulations than ABC. Furthermore, ML methods need not reexamine these simulations in order to perform downstream prediction, unlike ABC, and thus further inference is very fast.

A final difference between ML and ABC is that of interpretability. In the realm of ABC it is not clear which summaries are responsible for a signal. On the other hand many ML methods allow direct measurement of each feature’s contribution. Thus, despite their use of algorithmically generated models, ML algorithms are far from black boxes.

## Inferring Demography and Recombination

Another emerging use of supervised ML in population genetics has been for inference of demographic history and recombination rates. Indeed much attention in the field has been placed on developing methods for the inference of population size histories and patterns of population splitting and migration [71–75]. ABC methods are among the most popular for inferring demographic histories [68]. Interestingly, several groups have experimented with augmenting ABC by using ML for selecting the optimal combination of summary statistics [76] or even generating them [77]. While this is a promising direction for feature engineering, others have directly used ML to estimate posterior distributions of demographic parameters. For instance, Blum and Francois [70] used a feed-forward *artificial neural network* (ANN) to learn the mapping of summary statistics onto parameters with excellent results, particularly with respect to computational cost savings.

In addition to demographic parameter estimation, supervised ML has been used recently for demographic model selection (a possibility pointed to by Blum and Francois). For instance, Pudlo et al. [78] showed that random forests outperform ABC in both accuracy and computational cost when performing demographic model selection, along with greater robustness to the choice of summary statistics included in the input vector. In a recent preprint [79], we apply Extra-Trees classifiers to a problem of locus-specific demographic model selection: that of identifying regions with gene flow between a pair of closely related species with far greater accuracy than previous methods. Thus in general, ML methods show great promise in demographic estimation and model selection, and may soon be the preferred choice over ABC.

Supervised ML has also been applied to characterize rates and patterns of recombination in the genome. This work has again been done with or without simulation of training data. For instance Adrian et al. [80] trained a random forest classifier to distinguish among recombination rate classes on the basis of sequence motifs to show that such motifs are predictive of recombination rate in *Drosophila melanogaster*. This work used annotated rates of recombination based on a classical population genetics estimator to define the training set. On the other hand, Haipeng Li’s group has developed methodology [81, 82] that uses *boosting* to infer recombination rate maps from large sample sizes on the basis of simulated training data. Their latest method, FastEPRR, has much greater computational efficiency than and equal accuracy to the widely used LDhat [83]. Although application of supervised ML methods to this problem has begun only recently, the success of FastEPRR suggests the potential of future gains using these approaches.

## Co-estimation of selection and demography

It is well known that demographic events can mimic the effects of selection [54] and conversely that selection can confound demographic estimation [84, 85]. This implies that, although one can attempt to design more robust approaches (e.g. S/HIC, discussed above), the ideal strategy would be to simultaneously make inferences about both of these phenomena. How then can one perform co-estimation of parameters related to multiple evolutionary phenomena? A promising approach that utilizes supervised ML, in this case deep learning, was recently introduced by Sheehan and Song [17]. They developed a deep neural network, called evoNet, to simultaneously infer population size changes in a three-epoch model and detect selective sweeps. What makes this research particularly important is that Sheehan and Song performed simultaneous classification of loci into selective classes and demographic parameter estimation (based on averages estimated over loci classified as neutral), through the use of a neural network architecture that outputs both categorical and continuous parameters. This inherent flexibility of ML, and deep learning architectures in particular, opens up a whole slew of opportunities for doing population genomic inference in ways that have never before been possible (discussed below).

## Concluding Remarks and Future Directions

The future of population genomic analysis rests in our ability to make sense of large and ever growing datasets. Toward this end, supervised ML techniques represent a new paradigm for analysis, one uniquely suited for making inferences in the context of high-dimensional data produced by an unknown or imprecisely parameterized model. Here we have reviewed a selection of early applications of supervised ML tools to population genomic data. The overwhelming take-home is that supervised ML provides robust, computationally efficient inference for a number of problems that are difficult to gain traction on via classical statistical approaches.

We believe that population genetics is now poised for an explosion in the use of supervised ML approaches. Deep learning in particular, with its incredibly flexible input and output structure, should be an important area of future research, and its earliest application [17] has yielded the critical ability to co-estimate selection and demography, a central goal of population genetics analysis over the past 15 years. Indeed, deep learning could potentially alter the way that we even think about the nature of our input data itself. For example one flavor of deep learning, convolutional neural networks (CNNs), have made astounding advances in our ability to learn parameters from image data [86]. Rather than learning on population genetic summary statistics calculated from a multiple sequence alignment (e.g. [8, 16]), one could instead treat an image of the alignment itself as input. While these data would be extremely high dimensional, the structure of CNNs allows them to implicitly perform dimensionality reduction while capturing salient structures in the input data [87], allowing for accurate and efficient classification and regression (additional possible future avenues of ML in population genetics are listed in the **Outstanding questions** box). In general, the current explosion in deep learning research promises future improvements in our ability to make evolutionary inferences well beyond current capabilities; the challenge for population geneticists then is to adapt such methods for our own uses.

#### Outstanding questions

- While a few comparisons have shown that ML can outperform ABC, a more thorough assessment of the strengths and limitations of each approach across a variety of problems (e.g. on simulated data) is in order. In what scenarios would either strategy be preferable?
- Like more traditional methods, ML applications relying on simulated training data must make modeling assumptions. To what extent can ML methods be made more robust to these assumptions (e.g. by appropriately designing the feature vector as done by S/HIC, or through simulating a greater breadth of training examples)?
- ML methods have the ability to infer the values of multiple parameters simultaneously. How feasible will parameter estimation be in more complex evolutionary models using ML tools such as deep neural networks?
- As described here, supervised ML relies on summaries of population genetic data as feature vectors, but what summaries are best and can we do better than standard population genetic statistics? The recent rise of convolutional neural networks for image recognition suggests that encoding alignments as images might enable more powerful population genetic inference—how best can we encode population genetic data?
- A type of ANN called generative adversarial networks has been shown to generate data examples that can mimic true data with increasing accuracy. Can such methods be used as a substitute for population genetic simulation, perhaps to generate very large samples and chromosomes that are computationally costly to simulate?
- Applications of supervised ML to population genetic data can be quite involved, necessitating simulating data, encoding both simulated and real data as feature vectors, training the algorithm, and applying it. Can efforts to create self-contained, efficient, and user-friendly software packages capable of performing this entire workflow streamline this approach and make it more accessible to researchers?
- While point estimation of population genetic model parameters is important, equally important is establishing credible intervals on our parameter estimates. How can we most effectively use ML for estimating intervals associated with parameter estimates?

## Acknowledgments

We thank Alexander Xue, Matt Hahn, Parul Johri, Peter Ralph, and Yun Song’s group for comments on this manuscript. We also thank Justin Blumenstiel and Lex Flagel for discussions about image classification in population genetics. DRS was supported by NIH award no. K99HG008696. ADK was supported by NIH award no. R01GM117241.

## Glossary

Feature vector: A multidimensional representation of a data point made up of measurements (or features) taken from it (e.g. a vector of population genetic summary statistics measured in a genomic region).
Training: The process of algorithmically generating from a training set a function that seeks to correctly predict a datum's response variable by examining its feature vector.
Labeled data: Data examples for which the true response value (or label) is known.
Training set: A set of labeled examples for use during training.
Test set: A set of labeled examples for use during testing that is independent of the training set.
Loss function: A measure of how correctly an example's response variable was predicted.
Risk function: A measure of aggregated loss across an entire training set (e.g. the expected value of the loss function). We wish to minimize the value of the risk function during training.
Regression: A machine learning task where the value to be predicted for each example is a continuous number.
Classification: A machine learning task where the value to be predicted for each example is a categorical label.
Binary classification: A classification task in which there are two possible class labels, often termed positives and negatives.
Precision: In binary classification, the fraction of all examples classified as positives that are true positives (i.e. the number of true positives divided by the sum of the number of true positives and number of false positives). Also known as the positive predictive value.
Recall: In binary classification, the fraction of all positives that are correctly predicted as such (i.e. the number of true positives divided by the sum of the number of true positives and number of false negatives). Also known as sensitivity.
Confusion matrix: A table for visualizing accuracy in multi-class classification, which is simply the contingency table of the true and predicted classes for each example in a test set (see Figure Box 1 for an example).
Overfitting: When a model has achieved excellent accuracy in training data set but does not generalize well--i.e. the model has been tuned to precisely recognize patterns of noise in this set that are unlikely to be present in an independent test set. Sometimes referred to as overtraining.
*n*-fold cross validation: When only a small set of labeled data are available, cross validation can be used to measure accuracy. This process partitions the labeled data into *n* non-overlapping equally sized sets, and trains the predictor on the union of *n*-1 of these before testing on the remaining set. This is repeated *n* times, so that each of the *n* sets is used as the test set exactly once, and the average accuracy is recorded.
Boosting: A class machine learning techniques that seek to iteratively construct a set of predictors, weighing each predictor's influence on the final prediction according to its individual accuracy. Additionally, in most algorithms the new predictor to be added to the set focuses on examples that the current set of predictors has struggled with.
Support vector machine (SVM): A machine learning approach that seeks to find the hyperplane that optimally separates two classes of training data. These data are often mapped to highdimensional space using a kernel function. Variations of this approach can be performed to accomplish multi-class classification or regression.
Decision tree: A hierarchical structure that predicts an example's response variable by examining a feature, and branching to the right subtree if the value of that feature is greater than some threshold, and branching to the left otherwise. At the next level of the tree, another feature is examined. The predicted value is determined by which leaf of the tree is reached at the end of this process.
Random forest: An ensemble of semi-randomly generated decision trees. An example is run through each tree in the forest, and these trees then vote to determine the predicted value. Random forests can perform both classification and regression.
Artificial neural network (ANN): A network of layers of one or more “neurons” which receive inputs from each neuron in the previous layer, and perform a linear combination on these inputs which is then passed through an activation function. The first layer is the input layer (i.e. the feature vector) and the last layer is the output layer, yielding the predicted responses. Intervening layers are referred to as “hidden” layers.
Deep learning: Learning using ANNs or similarly networked algorithmic models that contain multiple "hidden" layers between the input and output layers.

## References

1. Breiman, L. (2001) Statistical modeling: The two cultures (with comments and a rejoinder by the author). from: Statistical science 16, 199–231

2. Elyashiv, E. et al. (2016) A genomic map of the effects of linked selection in Drosophila. from: PLoS Genet. 12, e1006130

3. Hinton, G. et al. (2012) Deep neural networks for acoustic modeling in speech recognition: The shared views of four research groups. from: IEEE Signal Processing Magazine 29, 82–97

4. Sebastiani, F. (2002) Machine learning in automated text categorization. from: ACM computing from: surveys (CSUR) 34, 1–47

5. Krizhevsky, A. et al., Imagenet classification with deep convolutional neural networks, Advances in neural information processing systems, 2012, pp. 1097–1105

6. Angermueller, C. et al. (2016) Deep learning for computational biology. from: Mol. Syst. Biol. 12, 878

7. Byvatov, E. and Schneider, G. (2003) Support vector machine applications in bioinformatics. from: Appl. Bioinformatics 2, 67

8. Lin, K. et al. (2011) Distinguishing positive selection from neutral evolution: boosting the performance of summary statistics. from: Genetics 187, 229–244

9. Mailund, T. et al. (2011) Estimating divergence time and ancestral effective population size of Bornean and Sumatran orangutan subspecies using a coalescent hidden Markov model. from: PLoS Genet. 7, e1001319

10. Kern, A.D. and Haussler, D. (2010) A population genetic hidden Markov model for detecting genomic regions under selection. from: Mol. Biol. Evol. 27, 1673–1685

11. Boitard, S. et al. (2009) Detecting selective sweeps: a new approach based on hidden Markov models. from: Genetics 181, 1567–1578

12. Pavlidis, P. et al. (2010) Searching for footprints of positive selection in whole-genome SNP data from nonequilibrium populations. from: Genetics 185, 907–922

13. Pybus, M. et al. (2015) Hierarchical boosting: a machine-learning framework to detect and classify hard selective sweeps in human populations. from: Bioinformatics 31, 3946–3952

14. Ronen, R. et al. (2013) Learning natural selection from the site frequency spectrum. from: Genetics 195, 181–193

15. Schrider, D.R. and Kern, A.D. (2015) Inferring selective constraint from population genomic data suggests recent regulatory turnover in the human brain. from: Genome Biol. Evol. 7, 3511–3528

16. Schrider, D.R. and Kern, A.D. (2016) S/HIC: Robust Identification of Soft and Hard Sweeps Using Machine Learning. from: PLoS Genet. 12, e1005928

17. Sheehan, S. and Song, Y.S. (2016) Deep learning for population genetic inference. from: PLoS from: Comput. Biol. 12, e1004845

18. Amit, Y. and Geman, D. (1997) Shape quantization and recognition with randomized trees. from: Neural Comput. 9, 1545–1588

19. Chen, D. et al., Blessing of dimensionality: High-dimensional feature and its efficient compression for face verification, Proceedings of the IEEE Conference on Computer Vision and Pattern Recognition, 2013, pp. 3025–3032

20. Kotsiantis, S.B. et al., Supervised machine learning: A review of classification techniques, 2007,

21. Ghahramani, Z. (2004) Unsupervised learning. In Advanced lectures on machine learning, pp. 72–112, Springer

22. Novembre, J. et al. (2008) Genes mirror geography within Europe. from: Nature 456, 98

23. Rabiner, L.R. (1989) A tutorial on hidden Markov models and selected applications in speech recognition. from: Proceedings of the IEEE 77, 257–286

24. Felsenstein, J. and Churchill, G.A. (1996) A Hidden Markov Model approach to variation among sites in rate of evolution. from: Mol. Biol. Evol. 13, 93–104

25. Siepel, A. et al. (2005) Evolutionarily conserved elements in vertebrate, insect, worm, and yeast genomes. from: Genome Res. 15, 1034–1050

26. Dutheil, J.Y. et al. (2009) Ancestral population genomics: the coalescent hidden Markov model approach. from: Genetics 183, 259–274

27. Hobolth, A. et al. (2007) Genomic relationships and speciation times of human, chimpanzee, and gorilla inferred from a coalescent hidden Markov model. from: PLoS Genet. 3, e7

28. Boitard, S. et al. (2012) Detecting selective sweeps from pooled next-generation sequencing samples. from: Mol. Biol. Evol. 29, 2177–2186

29. Cortes, C. and Vapnik, V. (1995) Support-vector networks. from: Machine Learning 20, 273–297

30. Quinlan, J.R. (1986) Induction of decision trees. from: Machine Learning 1, 81–106

31. Breiman, L. (2001) Random forests. from: Machine Learning 45, 5–32

32. Schapire, R.E. (1990) The strength of weak learnability. from: Machine Learning 5, 197–227

33. Bishop, C.M. (1995) Neural networks for pattern recognition, Oxford university press

34. LeCun, Y. et al. (2015) Deep learning. from: Nature 521, 436–444

35. Geurts, P. et al. (2006) Extremely randomized trees. from: Machine Learning 63, 3–42

36. Fay, J.C. and Wu, C.-I. (2000) Hitchhiking under positive Darwinian selection. from: Genetics 155, 1405–1413

37. Watterson, G. (1975) On the number of segregating sites in genetical models without recombination. from: Theor. Popul. Biol. 7, 256–276

38. Hudson, R.R. (2002) Generating samples under a Wright-Fisher neutral model of genetic variation. from: Bioinformatics 18, 337–338

39. Pedregosa, F. et al. (2011) Scikit-learn: Machine learning in Python. from: Journal of Machine from: Learning Research 12, 2825–2830

40. Liang, P. and Jordan, M.I., An asymptotic analysis of generative, discriminative, and pseudolikelihood estimators, Proceedings of the 25th international conference on Machine learning, ACM, 2008, pp. 584–591

41. Anderson, J. et al., The more, the merrier: the blessing of dimensionality for learning large gaussian mixtures, Conference on Learning Theory, 2014, pp. 1135–1164

42. Bengio, Y. and LeCun, Y. (2007) Scaling learning algorithms towards AI. from: Large-scale kernel from: machines 34, 1–41

43. Poggio, T. et al. (2017) Why and when can deep-but not shallow-networks avoid the curse of dimensionality: A review. from: International Journal of Automation and Computing, 1–17

44. Fu, Y.-X. and Li, W.-H. (1993) Statistical tests of neutrality of mutations. from: Genetics 133, 693–709

45. Fu, Y.-X. (1997) Statistical tests of neutrality of mutations against population growth, hitchhiking and background selection. from: Genetics 147, 915–925

46. Kelly, J.K. (1997) A test of neutrality based on interlocus associations. from: Genetics 146, 1197–1206

47. Kim, Y. and Nielsen, R. (2004) Linkage disequilibrium as a signature of selective sweeps. from: Genetics 167, 1513–1524

48. Tajima, F. (1989) Statistical method for testing the neutral mutation hypothesis by DNA polymorphism. from: Genetics 123, 585–595

49. Voight, B.F. et al. (2006) A map of recent positive selection in the human genome. from: PLoS from: Biol. 4, e72

50. Bengio, Y. (2009) Learning deep architectures for AI. from: Foundations and trends^®^ in Machine from: Learning 2, 1–127

51. Beaumont, M.A. et al. (2002) Approximate Bayesian computation in population genetics. from: Genetics 162, 2025–2035

52. Dunham, I. et al. (2012) An integrated encyclopedia of DNA elements in the human genome. from: Nature 489, 57–74

53. Altshuler, D.M. et al. (2012) An integrated map of genetic variation from 1,092 human genomes. from: Nature 491, 56–65

54. Simonsen, K.L. et al. (1995) Properties of statistical tests of neutrality for DNA polymorphism data. from: Genetics 141, 413–429

55. Meyer, K.A. et al. (2017) Differential gene expression in the human brain is associated with conserved, but not accelerated, noncoding sequences. from: Mol. Biol. Evol. 34, 1217–1229

56. Pollard, K.S. et al. (2006) Forces shaping the fastest evolving regions in the human genome. from: PLoS Genet. 2, e168

57. Maynard Smith, J. and Haigh, J. (1974) The hitch-hiking effect of a favourable gene. from: Genet. from: Res. 23, 23–35

58. Nielsen, R. et al. (2005) A scan for positively selected genes in the genomes of humans and chimpanzees. from: PLoS Biol. 3, e170

59. Hermisson, J. and Pennings, P.S. (2005) Soft sweeps molecular population genetics of adaptation from standing genetic variation. from: Genetics 169, 2335–2352

60. Orr, H.A. and Betancourt, A.J. (2001) Haldane’s sieve and adaptation from the standing genetic variation. from: Genetics 157, 875–884

61. Schrider, D.R. et al. (2015) Soft shoulders ahead: spurious signatures of soft and partial selective sweeps result from linked hard sweeps. from: Genetics 200, 267–284

62. Jensen, J.D. et al. (2005) Distinguishing between selective sweeps and demography using DNA polymorphism data. from: Genetics 170, 1401–1410

63. Nielsen, R. et al. (2005) Genomic scans for selective sweeps using SNP data. from: Genome Res. 15, 1566–1575

64. Schrider, D.R. and Kern, A.D. (2017) Soft sweeps are the dominant mode of adaptation in the human genome. from: Mol. Biol. Evol. 34, 1863–1877

65. Nei, M. and Li, W.-H. (1979) Mathematical model for studying genetic variation in terms of restriction endonucleases. from: Proceedings of the National Academy of Sciences 76, 5269–5273

66. Garud, N.R. et al. (2015) Recent selective sweeps in North American Drosophila melanogaster show signatures of soft sweeps. from: PLoS Genet. 11, e1005004

67. Messer, P.W. and Petrov, D.A. (2013) Population genomics of rapid adaptation by soft selective sweeps. from: Trends in Ecology & Evolution 28, 659–669

68. Beaumont, M.A. (2010) Approximate Bayesian computation in evolution and ecology. from: Annual review of ecology, evolution, and systematics 41, 379–406

69. Blum, M.G. et al. (2013) A comparative review of dimension reduction methods in approximate Bayesian computation. from: Statistical Science 28, 189–208

70. Blum, M.G. and François, O. (2010) Non-linear regression models for Approximate Bayesian Computation. from: Statistics and Computing 20, 63–73

71. Gutenkunst, R.N. et al. (2009) Inferring the joint demographic history of multiple populations from multidimensional SNP frequency data. from: PLoS Genet. 5, e1000695

72. Hey, J. and Nielsen, R. (2007) Integration within the Felsenstein equation for improved Markov chain Monte Carlo methods in population genetics. from: Proceedings of the National Academy of Sciences 104, 2785–2790

73. Li, H. and Durbin, R. (2011) Inference of human population history from individual whole-genome sequences. from: Nature 475, 493–496

74. Liu, X. and Fu, Y.-X. (2015) Exploring population size changes using SNP frequency spectra. from: Nat. Genet. 47, 555–559

75. Sheehan, S. et al. (2013) Estimating variable effective population sizes from multiple genomes: a sequentially Markov conditional sampling distribution approach. from: Genetics 194, 647–662

76. Aeschbacher, S. et al. (2012) A novel approach for choosing summary statistics in approximate Bayesian computation. from: Genetics 192, 1027–1047

77. Jiang, B. et al. (2015) Learning summary statistic for approximate Bayesian computation via deep neural network. from: arXiv preprint arXiv:1510.02175

78. Pudlo, P. et al. (2016) Reliable ABC model choice via random forests. from: Bioinformatics 32, 859–866

79. Schrider, D. et al. (2017) Supervised machine learning reveals introgressed loci in the genomes of from: Drosophila simulans and from: D. sechellia. bioRxiv, doi: 10.1101/170670

80. Adrian, A.B. et al. (2016) Predictive models of recombination rate variation across the Drosophila melanogaster genome. from: Genome Biol. Evol. 8, 2597–2612

81. Gao, F. et al. (2016) New software for the fast estimation of population recombination rates (FastEPRR) in the genomic era. from: G3: Genes, Genomes, Genetics 6, 1563–1571

82. Lin, K. et al. (2013) A fast estimate for the population recombination rate based on regression. from: Genetics 194, 473–484

83. McVean, G.A. et al. (2004) The fine-scale structure of recombination rate variation in the human genome. from: Science 304, 581–584

84. Ewing, G.B. and Jensen, J.D. (2016) The consequences of not accounting for background selection in demographic inference. from: Mol. Ecol. 25, 135–141

85. Schrider, D.R. et al. (2016) Effects of Linked Selective Sweeps on Demographic Inference and Model Selection. from: Genetics 204, 1207–1223

86. Sermanet, P. et al. (2013) Overfeat: Integrated recognition, localization and detection using convolutional networks. from: arXiv preprint arXiv:1312.6229

87. Graham, B. (2014) Fractional max-pooling. from: arXiv preprint arXiv:1412.6071

